# Targeting Wnt Signaling and DNAJB6/MRJ-L as a Dual Anti-RSV Strategy: Insights into a Positive Regulatory Loop

**DOI:** 10.1101/2025.03.27.645670

**Authors:** Chun-Yi Lu, Peng-Yeh Lai, Jen-Min Huang, Luan-Yin Chang, Ting-Yu Yen, Woan-Yuh Tarn, Li-Min Huang

**Author notes:** Corresponding author: (LMH), (WYT). E-mail address: Chun-Yi Lu, Peng-Yeh Lai, Jen-Min Huang, Luan-Yin Chang, Ting-Yu Yen, Woan-Yuh Tarn, Li-Min Huang. Author contributions: Chun-Yi Lu: Validation, Writing – Original Draft Preparation, Writing – Review & Editing Peng-Yeh Lai: Investigation, Formal Analysis, Investigation, Methodology, Visualization, Writing – Original Draft Preparation, Writing – Review & Editing Jen-Min Huang: Validation, Writing – Review & Editing Luan-Yin Chang: Validation, Writing – Review & Editing Ting-Yu Yen: Validation, Writing – Review & Editing Woan-Yuh Tarn: Conceptualization, Methodology, Project Administration, Visualization, Writing – Original Draft Preparation, Writing – Review & Editing Li-Min Huang: Conceptualization, Supervision, Funding Acquisition, Writing – Original Draft Preparation, Writing – Review & Editing.

## Abstract

Respiratory syncytial virus (RSV) is a major cause of severe respiratory infections, yet effective treatments are lacking. We found that the molecular chaperon DNAJB6/MRJ plays an essential role in RSV replication. Depletion of the long isoform of MRJ (MRJ-L) suppresses RSV replication. Transcriptomic analysis revealed that MRJ-L depletion downregulates Wnt signaling pathways. A pharmacological inhibitor of Wnt signaling suppressed RSV propagation and unexpectedly reduced MRJ-L expression, suggesting a positive regulatory loop between Wnt signaling and MRJ-L expression. Notably, simultaneous inhibition of Wnt signaling and MRJ-L additively suppressed RSV replication, suggesting that the Wnt-MRJ-L axis may serve as a new therapeutic target. This study provides insights into host-RSV interactions and potential antiviral strategies.

**Author Summary:** The molecular chaperone DNAJB6/MRJ has been implicated in the replication of respiratory syncytial virus (RSV), although the precise mechanisms remain unclear. In this study, we discovered that MRJ may influence RSV replication via Wnt signaling pathways. Specifically, we demonstrated that Wnt signaling inhibitor Wnt-C59 significantly reduced RSV replication by suppressing the synthesis of viral mRNA and genome/antigenome. Moreover, a positive feedback loop of the Wnt-MRJ axis may play a critical role in regulating RSV replication. Importantly, RSV replication was suppressed additively by inhibition of Wnt signaling and depletion of MRJ-L. Thus, a dual-targeted therapeutic approach may be effective in combating RSV infections.

## Introduction

Respiratory syncytial virus (RSV) is the leading cause of acute lower respiratory tract infections in infants, young children, and older adults worldwide [1, 2]. While preventive measures like vaccines and monoclonal antibodies are emerging [3, 4], effective therapeutic options remain limited [5, 6]. There is an urgent need for more effective, affordable RSV therapies to reduce the global disease impact.

The human DnaJ/Hsp40 family member B6 (DNAJB6 or the mammalian relative of DnaJ, MRJ) has been implicated in a variety of viral infections, indicating MRJ as a potential multivalent drug target [7]. Human MRJ has two splice isoforms, MRJ-L and MRJ-S, that are generated by alternative use of terminal exons [8]. MRJ-L is involved in the infection and replication of human immunodeficient virus (HIV) and human cytomegalovirus (HCMV) through interacting with viral proteins [7, 9–12]. MRJ-L also facilitates RSV replication, but the underlying mechanism remains elusive [7].

Given the reduced replication of RSV in MRJ-L-depleted laryngeal epidermoid carcinoma HEp-2 cells [7], we investigated whether any cellular pathways are altered upon MRJ-L depletion, leading to suppression of RSV. This study revealed that MRJ-L depletion impacted the expression of Wnt signaling pathways. Wnt signaling is an evolutionarily conserved biological pathway crucial for embryonic development and maintaining adult tissue homeostasis [13]. Wnt signaling involves β-catenin-dependent (canonical) and β-catenin-independent (noncanonical) pathways [14, 15]. In the canonical pathway, Wnt binds to the Frizzled receptors, stabilizing and activating β-catenin in gene regulation [16]. Activation of the non-canonical pathway leads to actin polymerization or c-Jun-dependent transcription [17]. Interestingly, it has been reported that MRJ modulates Wnt/β-catenin signaling via different mechanisms. MRJ induces β-catenin degradation by upregulating the Wnt inhibitor dickkopf 1 (DKK1) via the transcription factor MSX1 or activating glycogen synthase kinase 3β (GSK3β) via protein phosphatase PP2A in cancer [18–21]. Moreover, Wnt signaling regulates host responses to a variety of DNA viruses, including herpes simplex virus [22], Epstein-Barr virus [23], HCMV [24], hepatitis virus type B [25], and RNA viruses including HIV [26, 27], hepatitis virus type C [28], dengue virus [29], Rift Valley Fever virus [30], Sendai virus [31], influenza virus[32] and severe acute respiratory syndrome coronavirus 2 [33]. Notably, RSV infection of human lung epithelial A549 cells results in β-catenin protein stabilization and activation [34]. Therefore, understanding the reciprocal relationship between viral infection and Wnt pathways is crucial.

Our finding that MRJ-L depletion influenced the expression of a large set of Wnt components in RSV permissive HEp-2 cells prompted us to investigate whether Wnt signaling impacts RSV replication and whether targeting this pathway can be an anti-RSV strategy.

## Results

### *MRJ-L* knockout affects Wnt signaling

Our previous study showed that MRJ-L depletion impeded RSV propagation [7], but its underlying mechanism remained to be elucidated. To understand how MRJ-L influences RSV replication, we examined whether any cellular pathway is altered upon MRJ-L depletion. *MRJ-L* knockout (KO) HEp-2 cells and the parental line were subjected to RNA-seq analysis. Gene set enrichment analysis (GSEA) using the curated gene sets from the molecular signatures database (MSigDB) identified the gene sets enriched upon *MRJ-L* knockout (adjusted *p*-value <0.25) (S2 Table). Seven of the 341 gene sets enriched upon *MRJ-L* knockout were related to Wnt signaling (S2 Table, yellow parts). The enrichment plot shows that Wnt ligand biogenesis and trafficking and Wnt signaling in kidney disease were generally downregulated (Fig 1A, upper and lower panels). Heatmap analysis revealed that most Wnt ligands and Frizzled receptors (FZDs) were downregulated, whereas several negative Wnt regulators, including Dishevelled binding antagonist of β-catenin 1 (*DACT1*), CXXC finger protein 4 (*CXXC4*), axis inhibition protein 2 (*AXIN2*), and Dickkopf 3 (*DKK3*) were upregulated (Fig 1B, S2 Table). All downregulated Wnt ligands act in the non-canonical pathway [35–38], although some may also function in the canonical pathway [39, 40]. Next, we verified several targets with a significant and consistent change (high *p*-value) in *MRJ-L*-knockout HEp-2 cells, including three Wnt ligands (*WNT6*, *WNT10A*, and *WNT11*) and three negative regulators (*DACT1*, *CXXC4*, and *Axin2*). RT-qPCR analysis revealed the expected change in the selected Wnt ligands (Fig 1C) and negative regulators (Fig 1D). Therefore, MRJ-L knockout downregulated several Wnt ligands and upregulated negative regulators of Wnt-β-catenin signaling. Nevertheless, the Heatmap still revealed upregulation of the Wnt ligand *WNT2B*, receptor *FZD2* and activator R-Spondin 3 (*RSPO3*) (Fig 1B). Intriguingly, β-catenin mRNA (*CTNNB1*) was minimally upregulated in *MRJ-L* knockout HEp-2 cells (Foldchange: 1.637, *p*-value: 8.04×10^-5^). Moreover, previous reports have indicated that MRJ-L overexpression reduces β-catenin expression [19, 20]. These complex results may result from differential expression of Wnt components, crosstalk between different Wnt pathways [41], different cell contexts, and overexpression or knockdown of MRJ-L.

**Fig 1.**
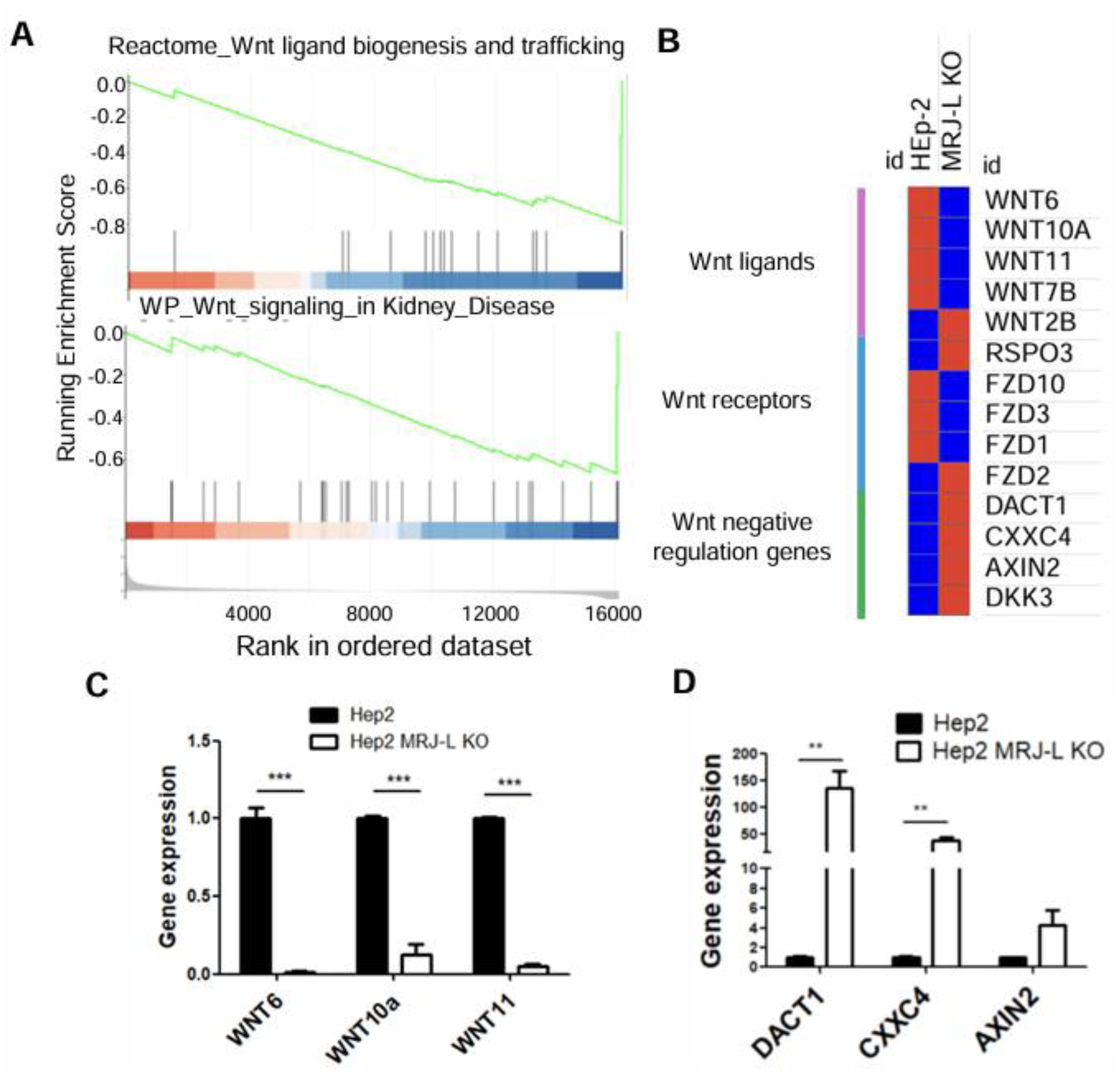
MRJ-L depletion reduces the expression of Wnt signaling factors. (A) GSEA enrichment plots show gene sets that were up/down-regulated upon *MRJ-L* knockout in HEp-2 cells. Upper panel: Wnt ligand biogenesis and trafficking genes. Lower panel: Wnt signaling in kidney disease. Enrichment score: −0.798 and −0.667; normalized enrichment score: - 1.929 and −1.803, *p-*value: both< 0.05. (B) Heatmap of Wnt signal genes in HEp-2 and *MRJ-L* KO HEp-2 cells was drawn by Morpheus website (Morpheus (broadinstitute.org)) according to the DEGs analysis (normalized mean counts, *p* value≦0.05). RT-qPCR analysis for the expression of Wnt ligand genes (C) and Wnt negative regulation genes (D) in HEp-2 and *MRJ-L* KO HEp-2 cells. Bar graphs show relative gene expression of *MRJ-L* KO/control; mean values were obtained from two independent experiments (N=2, **p≦0.01, *** p≦0.001).

### Inhibition of Wnt signaling reduces RSV propagation

RSV propagation was significantly reduced in *MRJ-L*-KO cells [7], in which several Wnt ligands were greatly diminished (Fig 1). Therefore, it was interesting to examine RSV infection in Wnt-inhibited or depleted cells. Wnt-C59, a porcupine inhibitor that inhibits Wnt secretion [42] (Fig 2A), was first tested for its impact on RSV infection. The cytotoxicity of Wnt-C59 in HEp-2 cells was evaluated using a tetrazolium-based cell viability assay. Wnt-C59 showed no deleterious effects on HEp-2 cells at concentrations up to 20 µM (Fig 2B). Next, we tested the effect of Wnt-C59 concentrations ranging from 0.5 to 5 µM on RSV infection. Increasing the doses of Wnt-C59 gradually reduced the level of total viral proteins as assessed by Western blotting and RSV RNA detected by RT-qPCR using the primers complementary to the N protein region (Fig 2C and 2D, respectively). Moreover, a plaque assay showed that Wnt-C59 at a concentration of 2.5 or 5 µM potently reduced viral titers (Fig 2E). We have also examined the cytopathic effect (CPE) of RSV infection, *i.e.* syncytia formation by fusion between adjacent cells. Wnt-C59 significantly reduced RSV-induced CPE in a dose-dependent manner (Fig 2F), suggesting that Wnt-C59 exhibited multiple inhibitory effects on RSV infection, including reduced viral RNA and protein levels, viral titers, and CPE.

**Fig 2.**
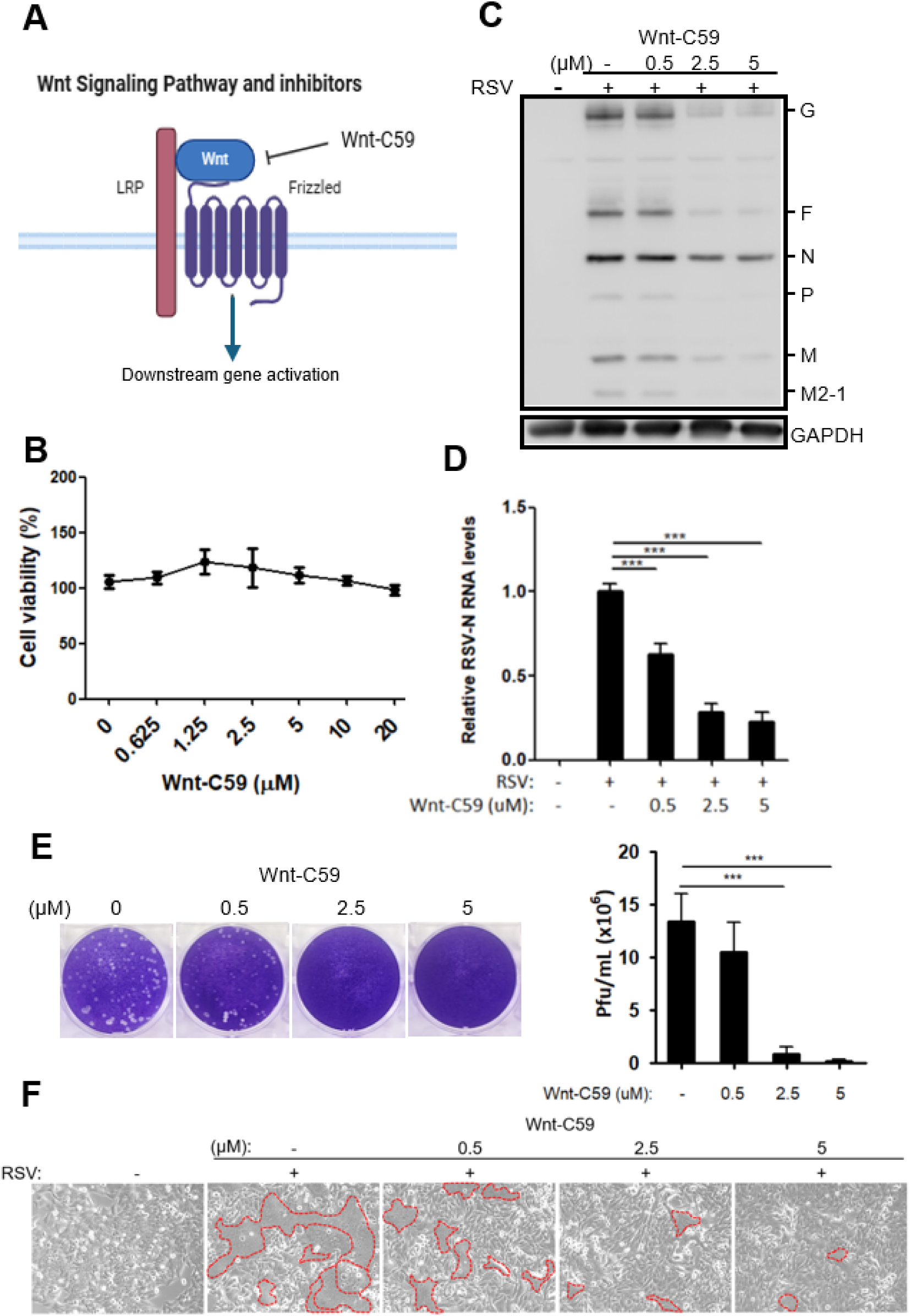
Wnt inhibition reduces RSV propagation. (A) The diagram shows Wnt/β-catenin signaling inhibitor, Wnt-C59 used in this study. The illustration was inspired by and created in BioRender. (B) Cytotoxicity of Wnt-C59 in HEp-2 cells. HEp-2 cells were seeded in 96 well plate and treated with different doses of Wnt-C59 for 48h. Cell viability was analyzed by using a tetrazolium-based MTT assay. Data was mean values obtained from two independent experiments. (C) HEp-2 cells were infected with RSV at an MOI of 0.05 followed by treatment with increasing doses of Wnt-C59 for 48h. Immunoblotting was performed using antibodies against RSV proteins and GAPDH. (D) The level of RSV RNA was determined by RT-qPCR and normalized to that of β-actin. The bar graph shows the relative level of RSV-N RNA in Wnt-C59-treated cells compared to mock-treated cells (N= 3). (E) The viral titer was determined by the plaque assay using culture supernatants. (F) As in panel C, cell morphologies were visualized by Nikon TS100 microscope. Red dashed line indicates syncytia; this experiment has been repeated. The bar graph shows the relative viral titer of Wnt-C59 treatment to mock-treatment (N= 3). **p≦0.01, *** p≦0.001.

### Wnt-C59 inhibits RSV mRNA expression and genome synthesis

To elucidate the mechanism of Wnt-C59 in combating RSV, we first examined the viral entrance step. HEp-2 cells were treated with Wnt-C59 before (pretreatment) or after (posttreatment) RSV infection (Fig 3 A). Pre-treatment had a similar effect on the expression of RSV proteins (Fig 3B) and N RNA to that of post-treatment (Fig 3C). This result suggested that Wnt-C59 hindered RSV replication in a step other than the viral entrance. Next, we examined the initial steps after RSV entry into host cells, *i.e.* mRNA transcription and RNA replication (Fig 3D). Wnt-C59 at 2.5 and 5 µM reduced the level of RSV N mRNA, antigenome and genome by 50-75% (Fig 3E), indicating that Wnt-C59 inhibits RSV mRNA expression and replication. Moreover, cells pretreated with Wnt-C59 prevented RSV infection, providing a viral prevention method.

**Fig 3.**
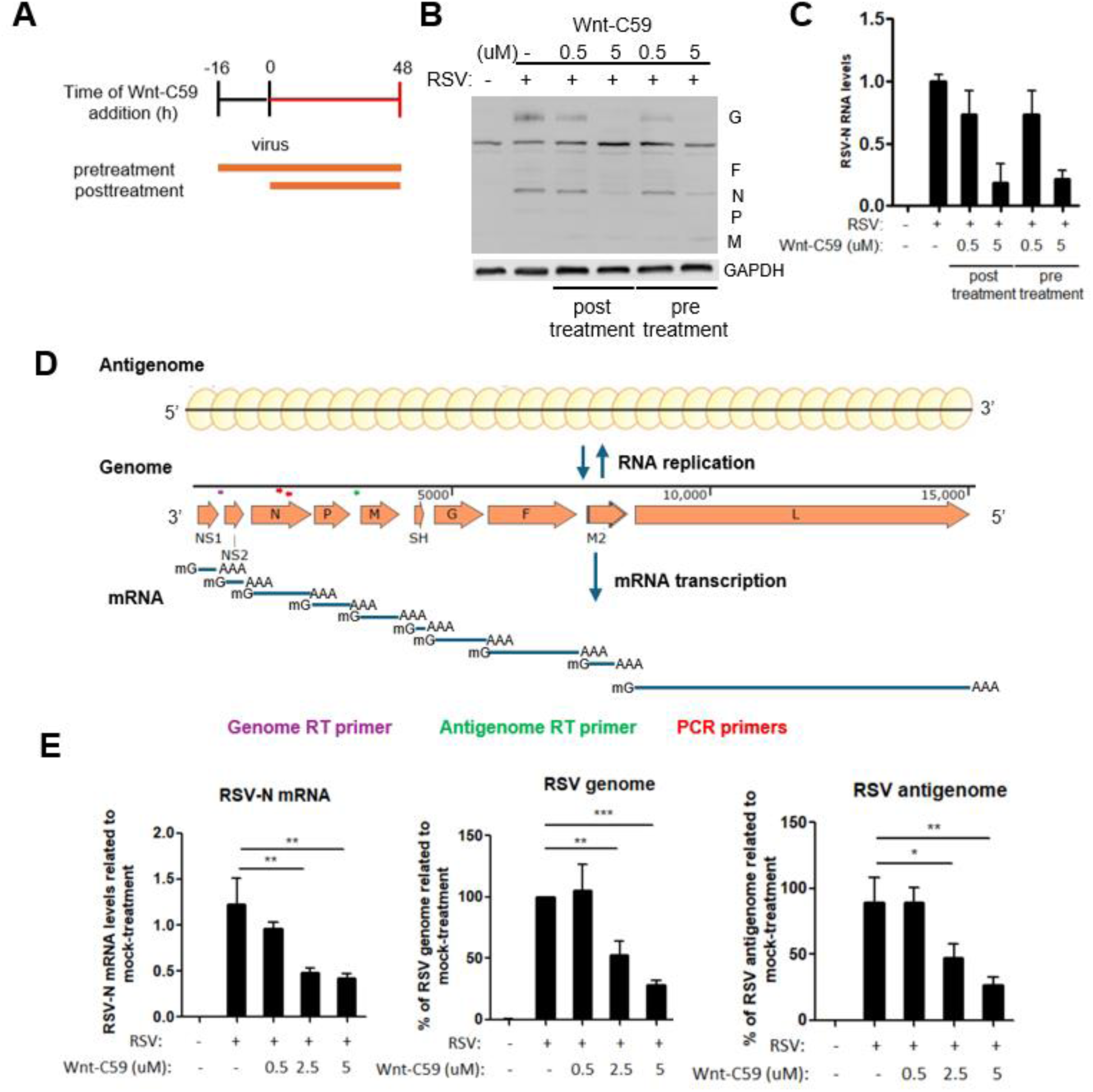
Wnt-C59 treatment reduces RSV mRNA expression and genome synthesis. (A) Schematic representation shows the time for Wnt-C59 addition versus virus infection. For post-treatment, HEp-2 cells were infected with RSV followed by treatment with Wnt-C59 for 48 h. For pre-treatment, HEp-2 cells were treated with Wnt-C59 for 16 h followed by RSV infection for another 48 h. For both conditions, MOI of 0.05 was used. Immunoblotting and viral RNA were determined as in (B) and (C), respectively. (D) Diagrams of RSV genome, antigenome and mRNAs. The primers sites are indicated by arrowhead in different colors (purple: genome reverse transcription primer; green: antigenome reverse transcription primer; red: PCR primers). (E) HEp-2 cells were infected with RSV at an MOI of 0.05 followed by treatment with increasing doses of Wnt-C59 for 48h. Bar graphs show the relative levels of RSV N mRNA, genome and antigenome in Wnt-C59-treated cells compared to mock-treated cells (N=3), respectively. For all bar graphs, mean values were obtained from two or three independent experiments. *p≦0.05, **p≦0.01, *** p≦0.001.

### Wnt-C59 reduces MRJ-L isoform expression by upregulating CstF64

We have previously reported that RSV infection depends on MRJ-L. Downregulation of the cleavage stimulation factor 64 (CstF64) during monocyte differentiation increases MRJ-L expression via alternative splicing coupled alternative polyadenylation [5] (Fig 4A). Unexpectedly, we observed that Wnt-C59 increased the level of CstF64 (Fig 4B). Accordingly, a reduction in MRJ-L mRNA and protein levels was observed with an increase in MRJ-S levels (Fig 4B for RT-PCR and immunoblotting, and 4C for quantification). Therefore, inhibition of Wnt signaling could result in splice isoform switching of MRJ by increasing CstF64. This result was not caused by RSV, since Wnt-59 exhibited the same effect in non-infected cells (Fig 4D and 4E). Together, our results revealed that the suppression of MRJ-L reduced Wnt signaling, resulting in further reductions of MRJ-L, suggesting a positive regulatory loop between Wnt and MRJ-L (Fig 4F). This potential Wnt-MRJ-L regulatory loop may amplify the effect of Wnt inhibitors on combating RSV. RSV prefers macrophages over monocytes [7]. Wnt signaling is activated during monocyte differentiation into macrophages [43]. Therefore, inhibiting Wnt appeared to mimic monocytes with higher levels of CstF64 and MRJ-S, making cells unfavorable for RSV infection (Fig 4G).

**Fig 4.**
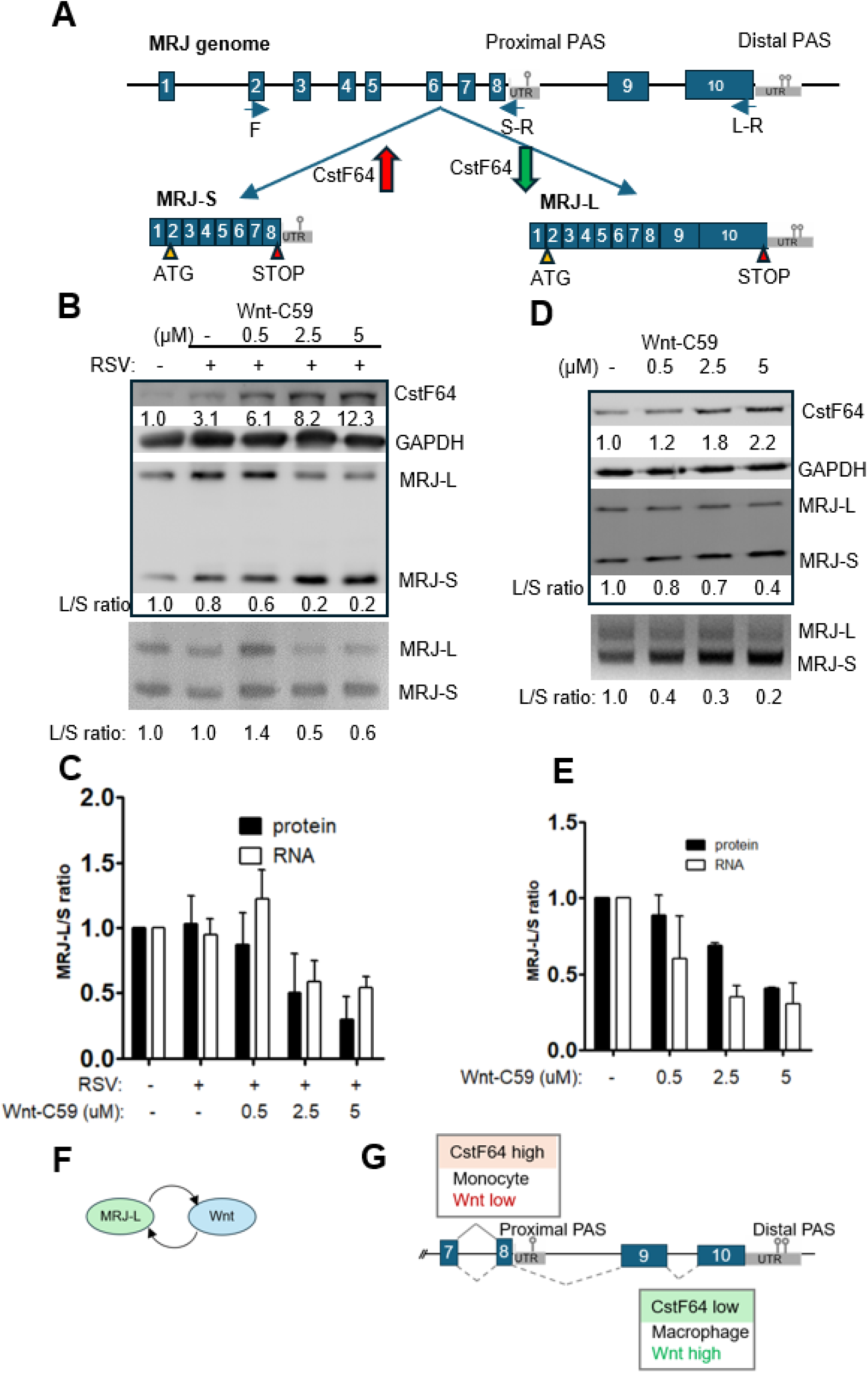
Wnt-C59 reduces MRJ-L expression by upregulating CstF64. (A) Schematic diagram showing the *MRJ* gene structure and two transcript isoforms generated by alternative splicing and polyadenylation. The level of CstF64 influences alternative 3’ end processing of the MRJ precursor mRNA. Arrows depict the primers used for RT-PCR of MRJ-L or MRJ-S. (B) HEp-2 cells were infected with RSV, followed by treatment with Wnt-C59 as shown in Figure 2C. Immunoblotting was performed using antibodies against MRJ, CstF64 and GAPDH (upper panel). RT-PCR was performed to detect MRJ-L and MRJ-S isoform mRNAs (lower panel). (C) Bar graph shows MRJ-L/MRJ-S protein and mRNA isoform ratios. (D) HEp-2 cells were treated with different concentrations of Wnt-C59 for 24 h. Immunoblotting was performed to detect CstF64, GAPDH and MRJ (upper panel). RT-PCR was performed to detect MRJ-L and MRJ-S isoform mRNAs (lower panel). (E) Bar graph shows MRJ-L/MRJ-S protein and mRNA isoform ratios. (F) A positive regulatory loop between MRJ-L and Wnt ligands. (G) MRJ-S expression is promoted by high levels of CstF64 in monocytes and during Wnt inhibition. MRJ-L expression is elevated when CstF64 levels are reduced in macrophages and in the presence of Wnt ligands.

### Inhibition of Wnt signaling and MRJ-L expression additively reduces RSV replication

In our previous study, MRJ-L morpholino oligonucleotide (MoMRJ) efficiently suppressed RSV replication [7]. Given that both MoMRJ and Wnt signal inhibitors effectively reduced RSV replication, we aimed to determine whether combining these two compounds could enhance anti-RSV activity. Our results revealed that MoMRJ and Wnt-C59, when applied separately, reduced both MRJ-L and RSV proteins (Fig 5A, compared lanes 8 and 9 to lane 7 for MoMRJ only, and lane 10 to lane 7 for Wnt-C59 only), viral N RNA expression (Fig 5B, green for MoMRJ only, and orange for Wnt-C59 only) and viral titer (Fig 5C, same as panel B). Notably, the combination treatment further reduced MRJ-L and viral gene expression and titers (Fig 5A, lanes 11 and 12, and Fig. 5B, 5C and 5D, red), indicating an additive effect between MoMRJ and Wnt-C59. These findings highlighted a potential combinatory approach in anti-RSV treatment by suppressing the MRJ-L-Wnt signaling pathway.

**Fig 5.**
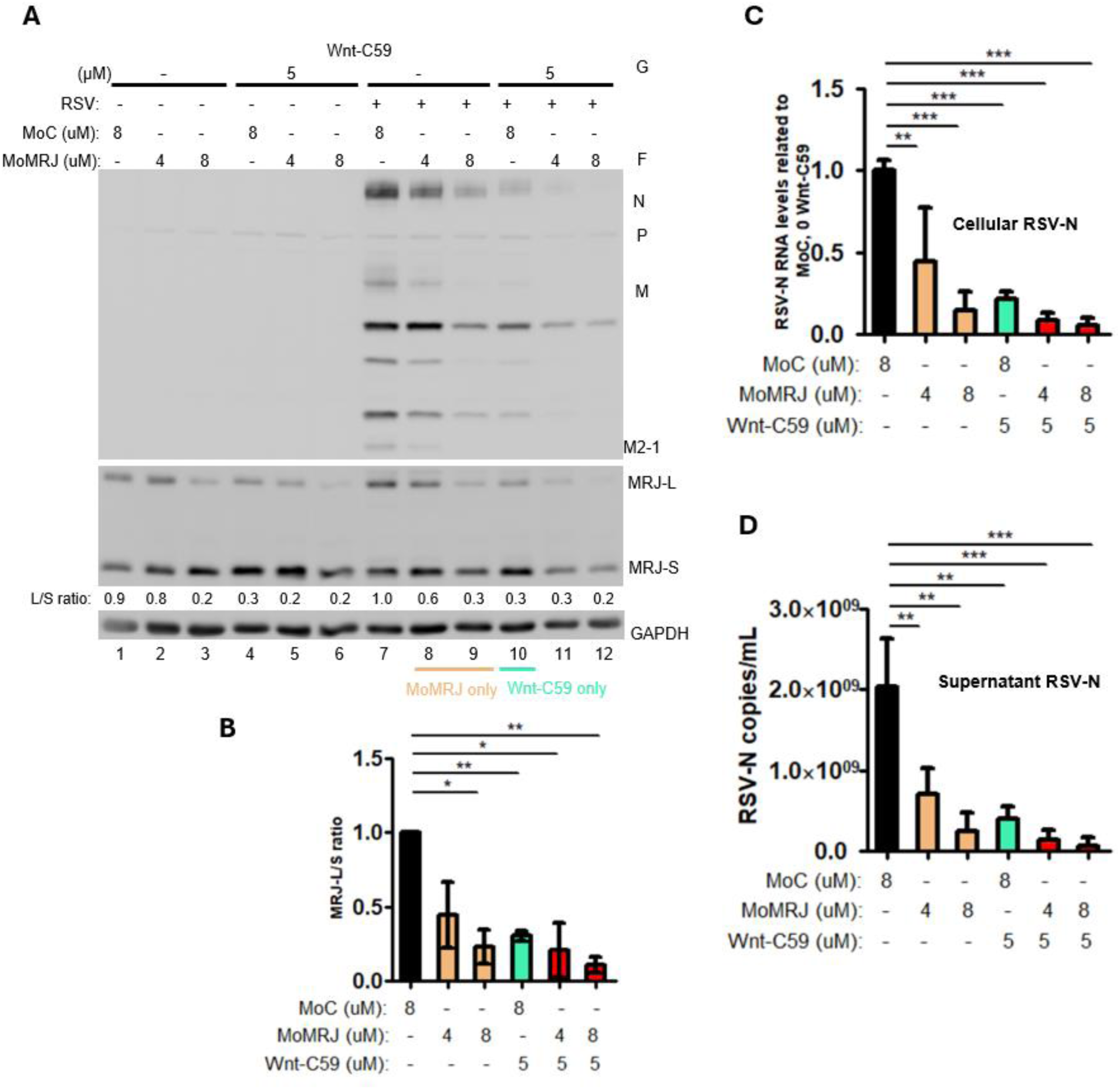
Wnt-C59 and MoMRJ additively reduce RSV replication. HEp-2 cells were treated with control (MoC) and/or MRJ morpholino oligonucleotide (MoMRJ) as indicated, followed by RSV infection in the absence or presence of 5 µM Wnt-C59. (A) Immunoblotting was performed using antibodies against RSV proteins, MRJ and GAPDH. The ratios of MRJ-L to MRJ-S are indicated below the blot and (B) is for quantification. For RSV infection samples, RT-qPCR was performed to detect RSV-N RNA in cells (C) and in the culture medium (D). For all bar graphs, mean values were obtained from two independent experiments.; *p≦0.05, **p≦0.01; *** p≦0.001.

## Discussion

RSV remains a significant health threat across all age groups. While preventive vaccines for pregnant women and the elderly and monoclonal antibodies for young children have recently become available, therapeutic options for RSV infection are still lacking. Some fusion and polymerase inhibitors have been evaluated in clinical trials, but none are currently clinically available. Our previous studies demonstrated that MRJ-L promotes RSV replication in the cells [7, 12]. This study advances our understanding of how MRJ-L facilitates RSV replication and suggests a new anti-RSV strategy.

### Wnt signaling is essential for RSV infection

MRJ-mediated Wnt regulation has been previously reported. MRJ-L upregulates Wnt signaling inhibitor DKK1 and prevents phosphorylation of GSK3β, thus promoting β-catenin degradation in certain cancer cells [18–20, 44], where it acts as a negative regulator of tumor growth and metastasis. We found that ablation of MRJ-L in HEp-2 cells downregulated several Wnt ligands and upregulated Wnt negative regulators. This contradiction may stem from context-specific regulatory mechanisms, which warrant further exploration. Notably, actin plays an important role in RSV endocytosis, replication, gene expression, and morphogenesis [45–49], and non-canonical Wnt ligands can regulate cell polarity and migration through actin polymerization [35, 38]. Our result that Wnt-C59 suppressed RSV-induced cytopathic effects (Fig 2) may echo the effect of non-canonical Wnt ligands in actin and microtubule cytoskeleton during RSV-induced cell fusion [50]. In addition, Wnt inhibition suppressed viral mRNA transcription and genome/antigenome synthesis (Fig 3), similar to the effects of MRJ-L depletion [7], suggesting the essential role of Wnt signaling in the life cycle of RSV.

### Wnt inhibitors as anti-viral agents

A variety of Wnt signaling inhibitors can inhibit viral replication. For example, KYA1797K and iCRT14 inhibit productive infection of HSV-1 in human and mouse neuronal cells and monkey kidney cells [51]. Moreover, iCRT14 also reduces titers of influenza virus, pseudorabies virus, and bovine viral diarrhea virus [32, 52, 53]. These inhibitors work differently in inhibiting viral infection and propagation: iCRT14 inhibits β-catenin-dependent transcription, whereas KYA1797K enhances the formation of the β-catenin destruction complex. Niclosamide, another Wnt/β-catenin inhibitor targeting Wnt co-receptor LRP6, suppresses RSV replication primarily through proapoptotic activity [54], leaving its role in Wnt-mediated RSV inhibition uncertain. Unlike the above inhibitors, Wnt-C59 functions to inhibit the secretion of both canonical and non-canonical Wnt ligands. We demonstrated that Wnt-C59 inhibited RSV replication in part by downregulating MRJ-L, emphasizing the potential for interfering with Wnt signaling in anti-RSV strategies.

### Dual targets for anti-RSV strategies

Our result unexpectedly revealed that Wnt inhibition affected MRJ isoform expression (Fig 4). Wnt signaling modulates RNA biogenesis at multiple levels, such as stabilizing COX2 mRNA via β-catenin and RNA-binding protein HuR interaction [55] or altering alternative splicing of genes like adenovirus E1A, Rac1 and SLC39A14 [56–58]. Here, Wnt inhibition affected MRJ alternative splicing and polyadenylation through CstF64. CstF64 expression changes during B cell activation and monocyte differentiation [7, 59] (Fig 4). Our previous results showed that MRJ-L levels rise during monocyte-to-macrophage maturation, a process where monocytes, naturally resistant HIV-1 [60, 61], become susceptible upon differentiation. While CstF64 generally affects alternative splicing and polyadenylation and may facilitate the maturation of primitive cells to more differentiated states, its decrease in macrophages correlates with MRJ-L expression [7]. Suppressing the Wnt pathway may represent an evolutionary mechanism to limit viral infections in eukaryotic cells, with MRJ-L playing a key role. How Wnt signaling modulates CsfF64 expression remains to be elucidated. Notably, MRJ-L expression and Wnt signaling appear to form a positive feedback loop that enhances RSV replication. Combining Wnt inhibition and MRJ-L suppression additively suppresses RSV replication (Fig 6), offering a potential novel dual-target anti-RSV strategy.

**Fig 6.**
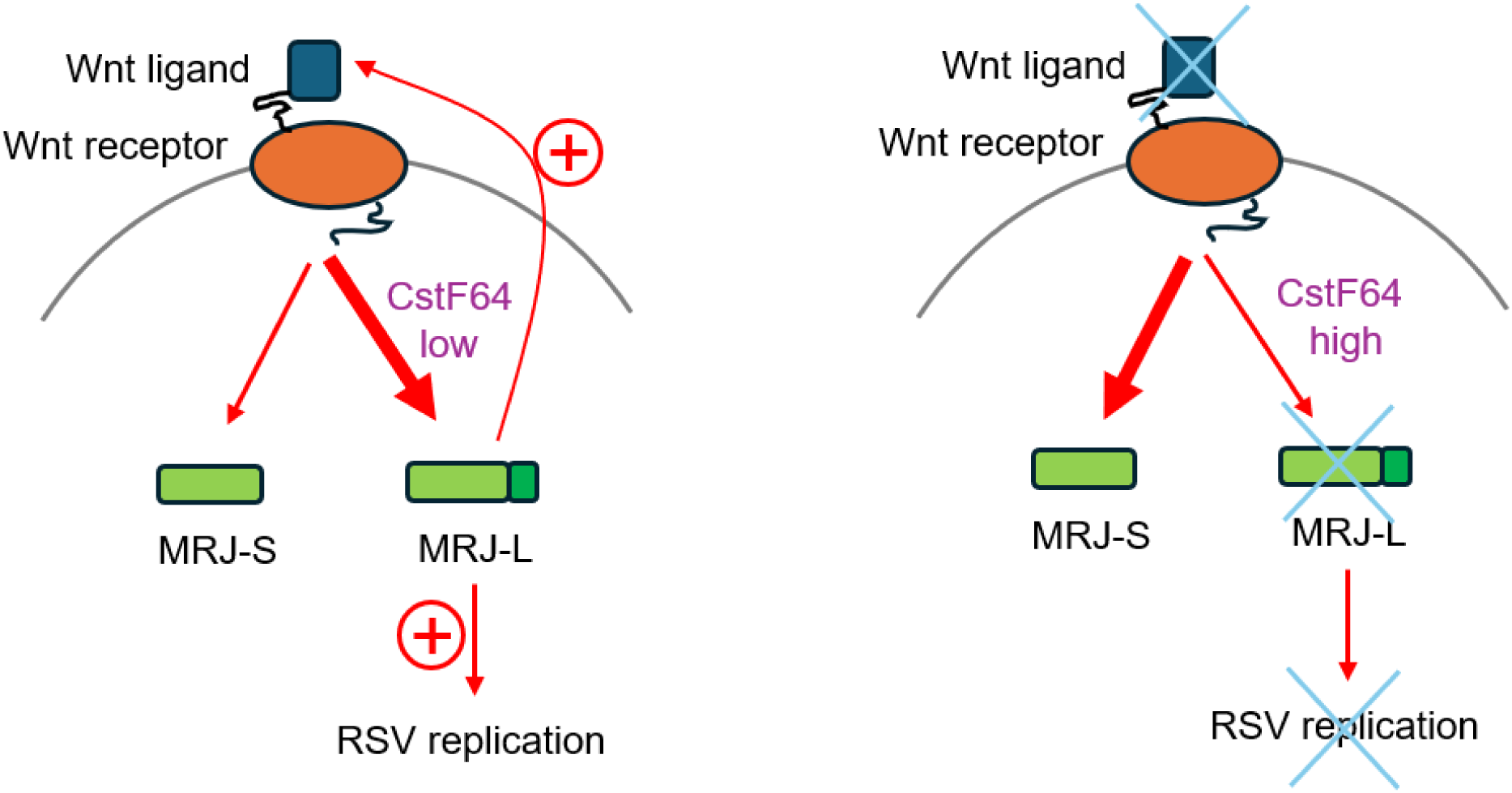
Wnt inhibition as an anti-RSV strategy. MRJ-L is crucial for RSV propagation. This study shows that Wnt inhibition suppressed RSV replication by upregulating CstF64, which in turn downregulated MRJ-L. Wnt signaling and MRJ-L establish a positive regulatory loop to facilitate RSV replication (Left). Wnt inhibition and MRJ-L suppression can additively suppress RSV replication, suggesting a new anti-RSV strategy (Right).

Our findings have broader implications for antiviral strategies beyond RSV. Given that Wnt signaling is dysregulated in infections by other RNA viruses (e.g., influenza, SARS-CoV-2) [32, 62], targeting the Wnt-MRJ-L axis may offer a multivalent approach for combating viral infections. Moreover, the additive effects of Wnt-C59 and MoMRJ suggest that combination therapies could enhance efficacy while minimizing resistance.

### Limitations

While our study highlights the Wnt-MRJ-L axis as a promising anti-RSV target, several limitations should be noted. First, our experiments were conducted in HEp-2 cells, which may not fully reflect RSV replication dynamics in primary human airway cells or *in vivo*. Second, the specificity of Wnt-C59 for RSV-related Wnt pathways remains unclear, as Wnt signaling is involved in many cellular processes and integrates with other signaling pathways. Future studies should validate these findings in more physiologically relevant models and explore potential off-target effects of Wnt inhibitors.

## Materials and methods

### Cell culture and viability assay

Human epithelial type 2 (HEp-2) (ATCC, CCL-23) cells were maintained in Dulbecco’s Modified Eagle Medium (DMEM) (Gibco), supplemented with 10 % fetal bovine serum (FBS) (Gibco) and antibiotics. Cells were maintained, and cell-based *in vitro* assays were performed in a humidified 37°C incubator with 5% CO_2_. Cell viability was measured using HEp-2 cells in 96-well plates. 10^4^ cells were seeded per well, followed by treatment with desired concentrations of Wnt signal inhibitor, Wnt-C59 (Abcam ab142216) in 2 % FBS-containing growth medium for 48 h. A tetrazolium-based MTT assay was performed following the manufacturer’s instructions (Sigma-Aldrich). Cells treated with DMSO were used as a positive control (100 % cell viability). The experiments were performed in duplicate.

### RSV propagation and infection

To propagate RSV, 2×10^6^ HEp-2 cells were seeded in a 10 cm dish for approximately 16 hours. When cell confluence reached 70-80%, cells were infected with RSV (A2 Strain, MOI 0.25) in DMEM medium for 2 h. Subsequently, 2% fetal bovine serum was supplemented. Virus-rich cell supernatant was harvested after 48-72 h when more than 50% of cells were detached from the cell culture flasks due to cytopathic effects. Cell debris was removed by centrifugation at 3200 ×g for 5 min. The supernatant was then aliquoted, and its viral titer was determined using a plaque assay.

### RT-PCR, RT-qPCR

Total RNA was extracted from cells using TRIzol Reagent (Invitrogen) and subjected to reverse transcription using random primers, oligo dT or gene-specific primers, and SuperScript III (Invitrogen) followed by PCR using gene-specific primers (Supplementary Table 1). PCR products were separated on 2% agarose gel. RT-qPCR was performed by LightCycler480 System (Roche Diagnostics).

### Immunoblot analysis

Cells were lysed in lysis buffer containing 20 mM Tris (pH7.5), 150 mM NaCl, 0.5 mM EDTA, 0.1% TritonX-100, and a protease inhibitors cocktail (Roche, 04-693-132-001). Proteins were separated on SDS-PAGE and transferred to nitrocellulose membranes. The subsequent steps were performed as previously described [12]. Antibodies targeting the following proteins or epitopes were used: RSV (Abcam, ab20745), MRJ (Abcam, ab198995), CstF64 (Abcam, ab72297), GAPDH (Proteintech, 10494-1-AP). Secondary antibodies included donkey anti-goat IgG (GeneTex, GTX232040), goat anti-mouse IgG (Jackson ImmunoResearch, 115-035-003), and goat anti-rabbit IgG (Jackson ImmunoResearch, 111-035-003). Detection was performed using an enhanced chemiluminescence detection kit (Thermo Fisher Scientific).

### RNA-seq and analysis

mRNA was enriched from total RNA using magnetic oligo-dT beads and fragmented with the KAPA Frag Kit (KAPA Biosystems, Switzerland) at 94°C for 4 minutes. First-strand cDNA was synthesized using random hexamer priming. Second-strand synthesis in the presence of dUTP and 3’ end A-tailing were combined, resulting in double-stranded cDNA with a dA protruding at the 3’ end. dsDNA adapters with 3’dT overhangs were ligated to the above cDNA. To select cDNA fragments of preferentially 300-400 bp in length, the library fragments were purified with the KAPA Pure Beads system (KAPA Biosystems). The library carrying appropriate adapter sequences at both ends was amplified using KAPA HiFi HotStart ReadyMix (KAPA Biosystems) along with library amplification primers. The strand marked with dUTP is not amplified, allowing strand-specific sequencing. At last, PCR products were purified using KAPA Pure Beads system, and the library quality was assessed on the Qsep 100 DNA/RNA Analyzer (BiOptic Inc., Taiwan). The original data obtained from high-throughput sequencing (Illumina NovaSeq 6000 platform) were transformed into raw sequenced reads by CASAVA base calling and stored in FASTQ format (Illumina). FastQC and MultiQC were used to check fastq files for quality. The obtained raw paired-end reads were filtered by Trimmomatic (v0.38) to eliminate low-quality reads, trim adaptor sequences, and eliminate poor-quality bases with the following parameters: LEADING:3 TRAILING:3 SLIDINGWINDOW:4:20 MINLEN:36. The obtained high-quality data (clean reads) was used for subsequent analysis. Read pairs from each sample were aligned to the reference genome (e.g., H. sapiens, GRCh38) by the HISAT2 software (v2.1.0). FeatureCounts (v2.0.0) was used to count the reads numbers mapped to individual genes. For gene expression, the “Relative Log Expression” (RLE) normalization was performed using DESeq2 (v1.26.0) with biological duplicates. Differentially expressed genes (DEGs) analysis was performed in R using DESeq2 (with biological replicates) based on Poisson distribution and negative binomial distribution models. The resulting *p*-values were adjusted using the Benjamini-Hochberg method to control the FDR. KEGG pathway enrichment analysis of DEGs was conducted using clusterProfiler (v3.14.3). Gene set enrichment analysis (GSEA) was performed with 1,000 permutations to identify enriched biological functions and activated pathways from the molecular signatures database (MSigDB) (Human), a collection of annotated gene sets for use with GSEA software [63, 64].

## Acknowledgments

The work was supported by the Ministry of Science and Technology, Taiwan (grant MOST 110-2314-B-002-097, MOST111-2314-B-002-026, MOST 112-2314-B-002-009 to L.-M.H).

---

**S1 Table. Primers used for detecting gene expression**

**S2 Table. Data of parental and *MRJ-L* KO HEp-2 cells Gene set enrichment analysis.**

